# Where is the boundary of the human pseudoautosomal region?

**DOI:** 10.1101/2024.06.10.598234

**Authors:** Daniel W. Bellott, Jennifer Hughes, Helen Skaletsky, Erik C. Owen, David C. Page

## Abstract

A recent publication describing the assembly of the Y chromosomes of 43 males was remarkable not only for its ambitious technical scope, but also for the startling suggestion that the boundary of the pseudoautosomal region 1 (PAR1), where the human X and Y chromosomes engage in crossing-over during male meiosis, lies 500 kb distal to its previously reported location^1^. Where is the boundary of the human PAR1? We first review the evidence that mapped the PAR boundary, or PAB, before the human genome draft sequence was produced, then examine post-genomic datasets for evidence of crossing-over between the X and Y, and lastly re-examine the sequence assemblies presented by Hallast and colleagues to see whether they support a more distal PAB. We find ample evidence of X-Y crossovers throughout the 500 kb in question, some as close as 246 bp to the previously reported PAB. Our new analyses, combined with previous studies over the past 40 years, provide overwhelming evidence to support the original position, and narrow the probable location of the PAB to a 201-bp window.

## Introduction

Ninety years ago, Koller and Darlington reported chiasmata between the rat X and Y chromosomes during male meiosis^2,3^. This led them to propose that the sex chromosomes would each consist of a “pairing segment” where X-Y crossing-over occurs, and a “differential non-pairing segment.” They proposed that while the differential non-pairing segment was completely sex-linked, the pairing segment should show partial sex-linkage, diminishing with genetic (recombinational) distance from the differential segments, and a total crossover frequency approaching 50 percent^3^. Nearly 50 years later, Burgoyne coined the term “pseudoautosomal” to describe this partial sex linkage in the pairing segment. We use the terms “pseudoautosomal region” (PAR) to refer to this region of partial sex linkage, and non-pseudoautosomal-Y (NPY) and non-pseudoautosomal-X (NPX) to refer to those regions that show complete sex-linkage.

Burgoyne posited that to ensure proper chromosome segregation, there was an obligatory crossover between each chromosome pair in male meiosis, and that for the human X and Y, this crossover would occur in a short arm region that was observed to pair under electron microscopy^4^. Further, he offered a set of criteria to identify a pseudoautosomal gene: in genetic mapping experiments, it should be partially linked to the X and Y, and not to an autosome; it should be physically mapped to the X and Y; and, because genes in the PAR would not require dosage compensation, genes in the PAR should show a strict dependence on sex chromosome dosage, in contrast to NPX genes subject to X-chromosome inactivation^4^. Only a few years later, in 1986, human *CD99* (formerly known as *MIC2*) was the first gene demonstrated to fulfill these criteria; it was physically mapped to the X and Y by deletion mapping, was not subject to X inactivation in human-rodent hybrid cells, and was observed to cross over with sex by pedigree analysis^5,6^.

The human X and Y chromosomes possess two PARs, one at the tip of the short arms and the other at the tip of the long arms (Figure 1a). PAR1, on the short arm (Xp and Yp), is all that remains of a larger ancestral eutherian PAR, while PAR2 is the result of a human-specific transposition from the tip of the long arm of the X chromosome to the Y chromosome^7^ (Figure 1a). The pseudoautosomal boundary (PAB) is the point on Xp and Yp where crossing-over between them ceases and the allelic PARs give way to divergent NPX and NPY sequences. The localization of *CD99* provided a foothold for further characterization of PAR1, and, in 1989, Ellis and colleagues fine mapped the PAB of PAR1, identifying an Alu repeat sequence inserted on the Y chromosome as the PAB^8^ (Figure 1b&c). This identification was supported by complementary genetic and physical mapping techniques. Pedigree-based genetic linkage maps, linkage-disequilibrium-based maps, and single sperm typing revealed patterns of crossing-over between X and Y chromosomes, while somatic cell hybrids, radiation-hybrid mapping, Southern blots, and clone-based sequencing all established physical linkage between the PAR1 and X– or Y-specific DNA sequences.

**Figure 1:**
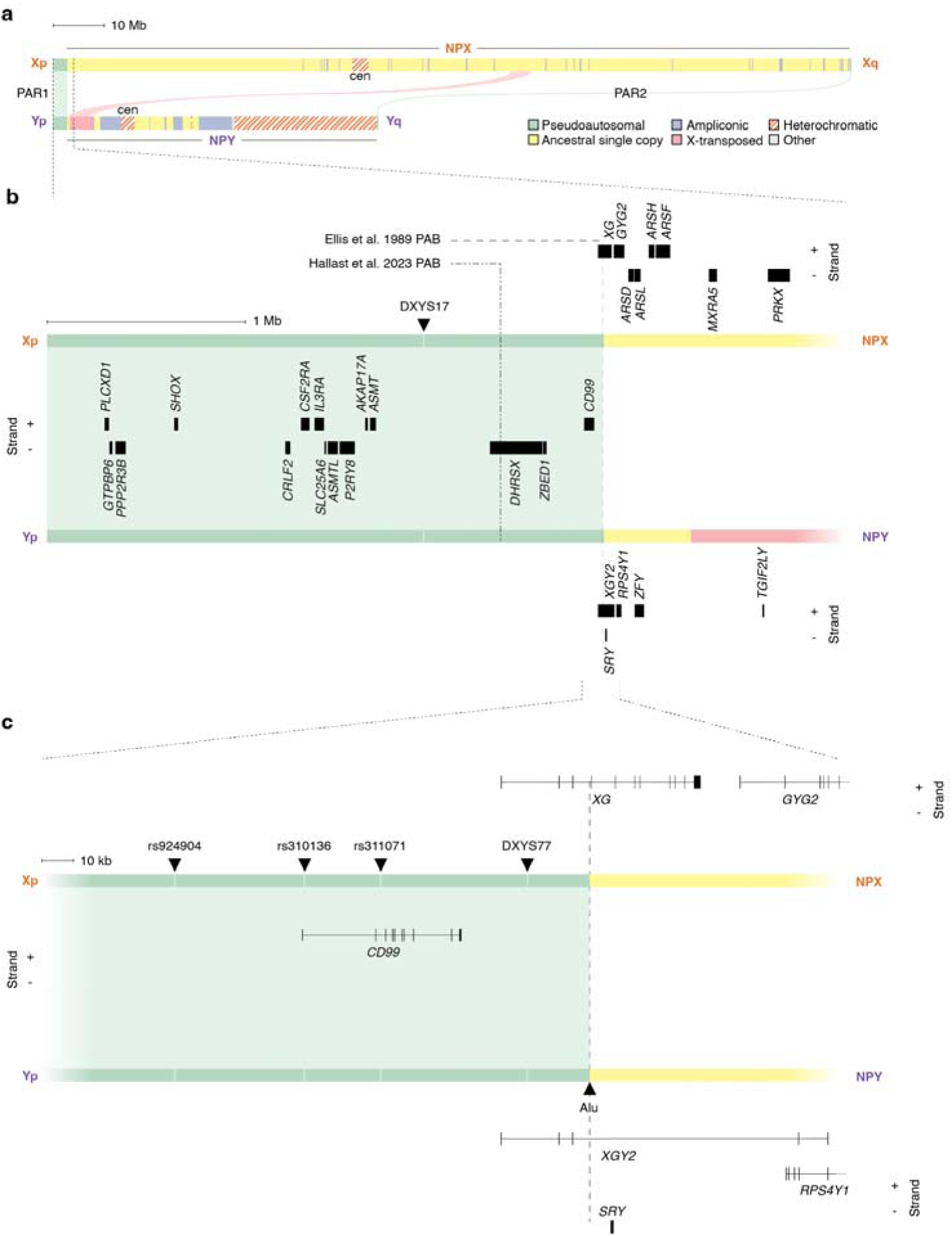
The boundary of human PAR1. Schematic representation of the human X and Y chromosomes, (a) showing the whole of each chromosome, including both PARs. Shading indicates four classes of euchromatic sequence, as well as heterochromatic sequences. Colored arcs indicate regions of >99% X-Y sequence identity. (b) Distal 4 Mb of the short arm of each chromosome, enlarged to show PAR1. Dashed lines: 2023 and 1989 PAB. Boxes: genes in PAR1, NPX, and NPY. Arrowheads: genetic markers. (c) 250 kb of each chromosome near the 1989 PAB. Genes shown as lines with arrowheads indicating direction of transcription, exons as boxes. *XG* and *XGY2* share three exons in the PAR.

Hallast and colleagues recently employed a novel approach to locate the PAB of PAR1, by using sequence variation in the human population as a proxy for crossing-over^1^. The recombination rate varies across the human genome, and regions of high recombination are observed to have a higher density of structural variants and indels. Both recombination and variation are highest in PAR1, followed by the autosomes, the NPX, and are lowest in the NPY. Hallast and colleagues observed that the 500-kb segment immediately distal to the 1989 PAB shows a low density of structural variants and indels among their sequenced Y chromosomes, and they reasoned that this observation is best explained by reduced or absent recombination in this interval. Hallast and colleagues concluded:

> “our data suggest that the boundary between the recombining pseudoautosomal region 1 and the non-recombining portions of the X and Y chromosomes lies 500 kb away from the currently established boundary”

This is indeed an arresting discovery – is it possible that the location of this critical landmark on the human sex chromosomes was mistaken, and the size of PAR1 has been overestimated by 20% for 35 years? What is the nature of the evidence for the 1989 PAB? First, we will review evidence accumulated over the past 40 years, and then we will present new analyses taking advantage of expanded genomic datasets, which all converge on confirming, and slightly refining, the original position of the PAB.

### Establishing the PAB

The earliest genetic maps of PAR1 were based on analysis of human pedigrees, using cloned DNA sequences as probes to identify restriction fragment length polymorphisms on Southern blots^6,9–13^. Rouyer and colleagues genotyped three pseudoautosomal markers across eight Centre d’Etude du Polymorphism Humain (CEPH) families to produce the first genetic linkage map of PAR1^11^. The map revealed a gradient of sex linkage from the most proximal to the most telomeric marker, and covered a distance of 50 cM (centiMorgans), indicating that, on average, one crossover occurs in PAR1 during each male meiosis^11^. However, *DXYS17*, the marker most closely linked to sex, was still 14 cM distal to the PAB (Figure 1b). Subsequently, Goodfellow and colleagues re-mapped these markers, along with the *CD99* gene, across an independent set of three pedigrees^6^. *CD99* was the proximal marker, mapping only 2.2 cM distal to biological sex^6^. Thus, the PAB for PAR1 must lie between *CD99* and the sex determining locus.

The tight linkage of *CD99* with sex inspired researchers to attempt to isolate the boundary between PAR1 and the NPY using *CD99* as a probe. Using a 49,XYYYY cell line as a highly enriched source of Y-linked DNA, Pritchard and colleagues used rare-cutting restriction enzymes, both singly and in pairs, to generate long restriction fragments which they resolved by pulsed-field gel electrophoresis^14^. They identified a 200-kb SfiI/ClaI fragment that hybridized to both *CD99* and NPY sequences^14^. Subsequently, Ellis and colleagues further refined this interval, beginning with a 110-kb restriction fragment containing their most proximal *CD99* and most distal NPY probes, and gradually narrowed this down to a 0.8-kb restriction fragment containing the boundary between PAR1 and NPY^8^. Armed with precise knowledge of the most proximal PAR sequences, they also isolated a 0.9-kb restriction fragment containing the boundary between PAR1 and NPX^8^. They sequenced both fragments and found they were 99% identical distal to a 303-bp Alu repeat insertion in the Y-derived sequence^8^. Proximal to the Alu insertion, X-Y identity drops to 77% for another 245 bases, and fails to align further^8^. They reasoned that the high X-Y identity in the sequence distal to the Alu insertion was maintained by crossing-over, and that the distal end of the Alu insertion marked the PAB (Figure 1c).

To establish that this sequence did constitute the PAB, it was critical to find additional evidence of crossing-over. Ellis and colleagues took a two-pronged approach, examining homologous X and Y sequences across catarrhine primates and sequencing this region from diverse human populations to identify polymorphisms^15,16^. Their evolutionary analysis found that while the Alu insertion was found across the Y chromosomes of great apes, it was absent from Old World monkeys^15^. However, even Old World monkeys maintained a crisp drop-off in X-Y identity between regions distal and proximal to the Alu insertion site^15^. This suggested that X-Y identity in the distal region was maintained by crossing-over since the split of Old World monkeys and great apes. By surveying 57 Y chromosomes and 60 X chromosomes from ten human populations, Ellis and colleagues identified single-nucleotide polymorphisms to infer patterns of crossing-over^16^. While there were fixed X-Y differences at positions 41 and 45 bases distal to the Alu insertion, they observed a site with a polymorphism shared by the X and the Y only 274 bases distal to the Alu^16^. Between these two sites, all Y chromosomes shared SNPs with a common X chromosome haplotype. This provided strong evidence that crossovers had occurred between 45 and 274 bases distal to the Alu insertion, localizing the human PAB to this 229-base interval.

Following the localization of the 1989 PAB, crossovers have repeatedly been observed in the 500 kb distal to this site by independent methods. Pedigree-based genetic maps rely on crossovers to infer the linear order and genetic distances among loci along a chromosome. In contrast, single-sperm typing isolates and genotypes single haploid sperm cells to reconstruct haplotypes without pedigree analysis and infer meiotic recombination events. Both methods have observed crossing-over between the 1989 PAB and markers only tens of kilobases away.

In producing the second iteration of the Rutgers Map of the human genome, Matise and colleagues mapped 28,121 markers on CEPH pedigrees, including three markers within *CD99* (rs924904, rs310136, and rs311071)^17^ (Figure 1c). These markers mapped from 5.12 to 2.79 cM distal to the 1989 PAB^17^. Flaquer and colleagues added more pseudoautosomal markers to the third iteration of the Rutgers map, and mapped two of these markers even closer to the PAB: rs310136, 87.6 kb and 0.7 cM away; and rs311071, 66.3 kb and 0.0007 cM distal to the PAB^18,19^. Subsequently, Hinch and colleagues generated a fine-scale map of the PAR; they observed 126 paternal crossovers in PAR1, of which at least 14 were in the 500 kb immediately distal to the 1989 PAB^20^. This region covers 5.1 cM in their map, and includes a recombination hotspot within the *XG* gene, which straddles the PAB on the X chromosome, but is disrupted on the Y chromosome^20^.

Schmitt and colleagues typed 903 flow-sorted sperm from two donors to map recombination among five loci in PAR1, and used 2,005 sperm from a third donor to map *DXYS77*, the marker closest to the 1989 PAB, at higher resolution^21^. *DXYS77* is located in an intron of the *XG* gene, 19.6 kb and 0.77 cM distal to the PAB (Figure 1c). Subsequent single-sperm typing analyses have not mapped PAR1 at such high resolution, but have also observed crossovers in the 500 kb distal to the 1989 PAB^22,23^.

In summary, several lines of evidence that revealed both historical and contemporary crossovers between X and Y chromosomes were used to support the 1989 PAB. Here we take advantage of a variety of publicly-available post-genomic datasets to re-examine this question in light of the more distal PAB proposed by Hallast and colleagues. We examine crossover events during primate evolution by aligning X and Y chromosomes from ape and Old World monkey genome assemblies. Similarly, we use sex differences in minor allele frequency in a genome-wide catalog of common human genetic variation to reveal historical patterns of crossing-over in the human population. Analysis of several independent single-sperm sequencing datasets allowed us to locate ongoing crossovers in the PAR of living individuals. Lastly, we use the known phylogenetic relationship between the NPY sequences in the genomes assembled by Hallast and colleagues to look for patterns of nucleotide substitution that are best explained by crossing-over. We find that all of these lines of evidence support the 1989 PAB.

## Methods

### Primate X-Y alignments

We downloaded primate X– and Y-linked sequences spanning the PAB from GenBank (Supplemental Table 1). Prior to alignment, we removed the Alu insertion at the 1989 PAB from ape Y-linked sequences, as well as an independent Alu insertion 300 bp proximal to the 1989 PAB in the colobus NPY. We used PRANK (v.170427)^24^ to generate a multiple alignment of sequences distal and proximal to the 1989 PAB (Supplemental Data 1). We used various windows of these multiple alignments to infer the phylogenetic relationships between sequences with PhyML (v.3.3.20190909)^25^ with 100 bootstrap replicates.

### Human polymorphisms from 1000 genomes

We downloaded phased individual-level variants from the PAR from the 1000 Genomes Project^26^. For each of 118,327 polymorphic sites in PAR1, we tabulated the number of males and females who were homozygous for the reference allele, heterozygous, and homozygous for the alternative allele (Supplemental Table 2). We used these counts to calculate sex-specific minor allele frequencies for males and females.

### Single sperm sequencing

We obtained raw sequence data dbGaP study phs001887 from the NCBI Sequence Read Archive. Using Bowtie2 (version 2.4.2)^27^, we aligned reads to a version of hg38 (GCF_000001405.40) with the chrY PAR masked. For each donor, we combined PAR alignments from all haploid cells using samtools (version 1.11)^28^, marked duplicate reads with sambamba (version 0.8.2)^29^, and called variants at sites in dbSNP with freebayes (version 1.3.2)^30^. We used vcffilter^31^ to restrict downstream analyses to biallelic SNPs with Q > 20, more than 2 reads each of reference and alternate alleles, and allele frequencies between 0.25 and 0.75. We identified X– and Y-bearing sperm by the depth of reads mapping to NPX and NPY sequences. In our analyses of the Sperm-seq data from phs001887, we also excluded reads with cell barcodes associated with cell doublets and bead doublets by Bell and colleagues^32^.

For each donor, we reconstructed X and Y haplotypes across PAR1 in two passes. In each pass we assigned the alleles at each variant site to either the X or Y donor haplotype to minimize the number of crossovers with more proximal sites, then, within each sample, localized crossovers between sites with variants assigned to different donor haplotypes. In the first pass, we flagged markers in tight double recombinants (closer than 278148 bases; that is, one tenth the length of PAR1, or ∼5 cM), as unreliable, and identified non-crossover samples. In the second pass, we excluded unreliable markers flagged in the first pass, as well as any markers with more than 10% discordant genotypes in non-crossover samples. We then removed any remaining tight double recombinants before calculating the local recombination rate.

### Reanalysis of Y chromosomes sequenced by Hallast and colleagues

Hallast and colleagues produced contiguous assemblies of the PAR-NPY boundary from ten samples (HG01890, HC02666, HC19384, NA19331, HG02011, HG03371, NA19317, HC18989, HG03009, and HG00358); we downloaded their assemblies (ftp://ftp.1000genomes.ebi.ac.uk/vol1/ftp/data_collections/HGSVC3/working/20230412_ sigY_assembly_data), and extracted the PAR contigs for further analysis.

To analyze clock-like evolution on the Y chromosome, we first masked simple repeats with RepeatMasker (version 4.1.5)^33^ and tandem repeats with trf (version 4.09)^34^, then aligned all ten Y chromosome assemblies with progressiveMauve (build date May 15 2023 at 08:46:41)^35^. We extracted 20-kb windows of the multiple alignment, excluding columns of gaps and masked characters, at 1-kb intervals. We used tree-puzzle (version 5.2)^36^ to conduct a likelihood ratio test to compare a model where sequences evolve using a single clock-like rate based on their relationships from the Y chromosome haplogroup tree, versus a model where rates are allowed to vary across branches. We obtained p-values from the test statistic reported by tree-puzzle (Supplemental Table 4), using the pchisq function and adjusted for multiple testing by Bonferroni correction with the p.adjust function in R (version 4.1.1)^37^.

To look for polymorphisms shared among X and Y chromosomes, we aligned homologous X and Y sequences from the ten genomes with contiguous PAR sequence and RP11-400O10, a human Y chromosome BAC spanning the 1989 PAB, and manually edited the resulting alignments in Unipro UGENE (version 49.1)^38^ (Supplemental Data 3)).

## Results

### Primate X-Y alignments

We took advantage of the rich genomic resources from multiple species to revisit the evolutionary analyses of Ellis and colleagues, using high-quality assemblies of primate X and Y chromosomes. Within the PAR, crossing-over will result in the DNA sequences of Y chromosomes being more similar to X chromosomes of the same species than to Y chromosomes of other species. Conversely, in the NPY, the absence of crossing-over will result in NPY sequences from different species being more similar to each other than to NPX sequences from the same species. We curated high-quality sequences adjacent to the 1989 PAB from the NPX and NPY from seven catarrhine primate species: human, chimpanzee, gorilla, orangutan, siamang, rhesus macaque, and colobus monkey (Supplemental Table 1, Supplemental Data 1). We constructed a multiple alignment from these sequences, and used these alignments to construct trees using a range of window sizes from the alignment, both distal and proximal to the 1989 PAB (Figure 2). Across all window sizes, even as narrow as 1 kb from the 1989 PAB (Figure 2d), we observe that sequences distal to the 1989 PAB group together by species, while sequences proximal to the 1989 PAB group together by chromosome. Examining this multiple alignment more closely, we find human X and Y chromosomes sharing an A>G substitution only 93 bases distal to the 1989 PAB (Figure 3). Since this substitution is also present on the chimpanzee Y, this suggests that it originated on the Y chromosome in the last common ancestor of humans and chimpanzees, and crossed over to the X chromosome in humans (Figure 3). The Alu element, from the cattarhine-specific AluY family^39^, was inserted into the Y chromosome after apes diverged from Old World monkeys, and has not crossed over to the X. This Alu insertion must have occurred at or near the ancestral catarrhine PAB; only 11 bases proximal to this site, all X chromosomes share a T where all Y chromosomes carry a G or A (Figure 3), confirming that crossing-over was suppressed in this proximal region before the divergence of apes and Old World monkeys.

**Figure 2:**
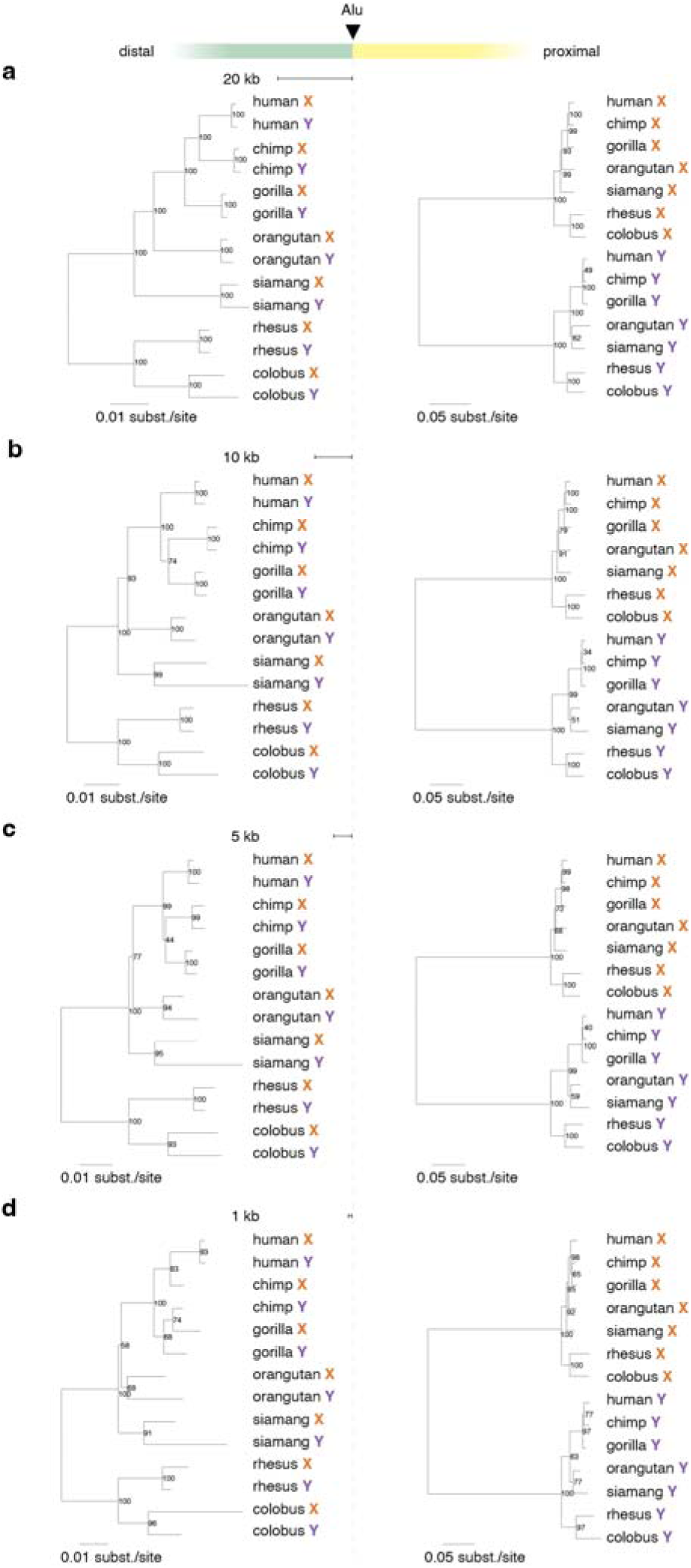
Phylogenetic relationships between primate X and Y sequences differ distal and proximal to 1989 PAB. Maximum likelihood trees based on multiple alignment of primate sex chromosome sequences in windows (a) 20 kb, (b) 10 kb, (c) 5 kb, or (d) 1 kb upstream and downstream of the 1989 PAB; nodes show support for branching order from 100 bootstrap replicates.

**Figure 3:**
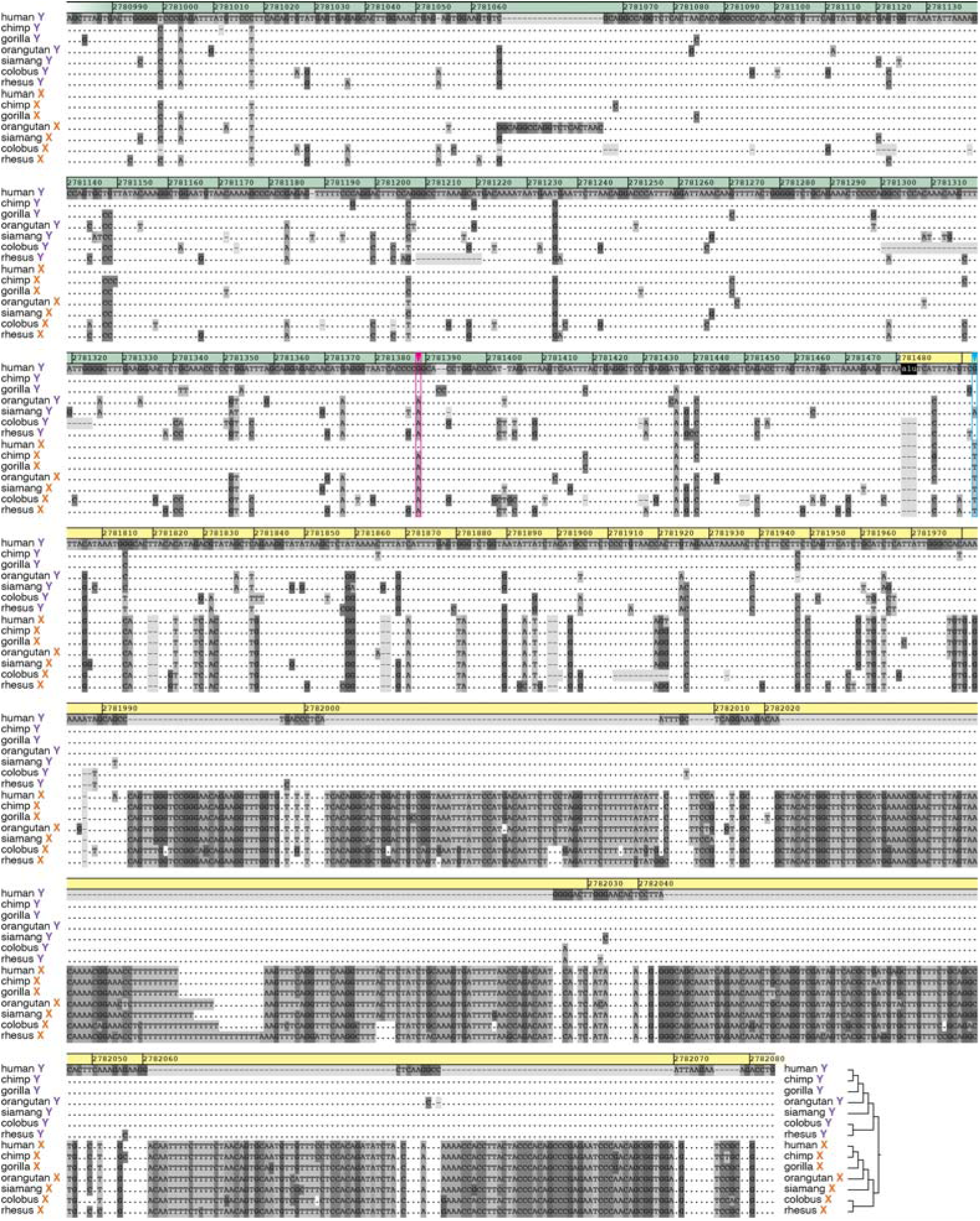
Multiple alignment of primate sex chromosomes shows sharp change in X-Y identity at the 1989 PAB. Multiple alignment of primate sex chromosome sequences showing the 1989 PAB at or near the ape-Y-specific Alu insertion (black). Dots indicate identity with human Y sequence; coordinates relative to hg38 reference. Tree indicates phylogenetic relationship in NPX and NPY regions. Magenta arrowhead and box marks crossover from Y to X in the human lineage; cyan arrowhead and box a fixed NPX/NPY difference predating the Alu insertion in apes.

### Human polymorphisms from 1000 Genomes

To gain additional insight into the history of crossing-over within the human population, we analyzed polymorphism data from the 1000 Genomes Project^26^. Strictly Y-linked regions with low or absent crossing-over with the X chromosome will accumulate mutations that are male-specific, leading to sex biases in the frequency of minor alleles. Using data from 1,599 males and 1,603 females, we calculated sex-specific minor allele frequencies for 118,327 polymorphic sites across PAR1 (Supplemental Table 2). We plotted the sex difference in minor allele frequency throughout PAR1 and observed that the differences between male and female minor allele frequencies are typically small, even in the 500 kb immediately distal to the 1989 PAB (Figure 4a). Within 10 kb of the 1989 PAB, alleles are more sex-biased, and this sex bias in minor allele frequencies increases rapidly as one moves proximally, nearing the maximum possible value of 0.5 (Figure 4a). Distal to this 10-kb segment adjacent to the 1989 PAB, male and female minor allele frequencies closely mirror each other, indicating that crossing-over has intermixed and homogenized the sets of alleles on the X and Y chromosomes (Figure 4b).

**Figure 4:**
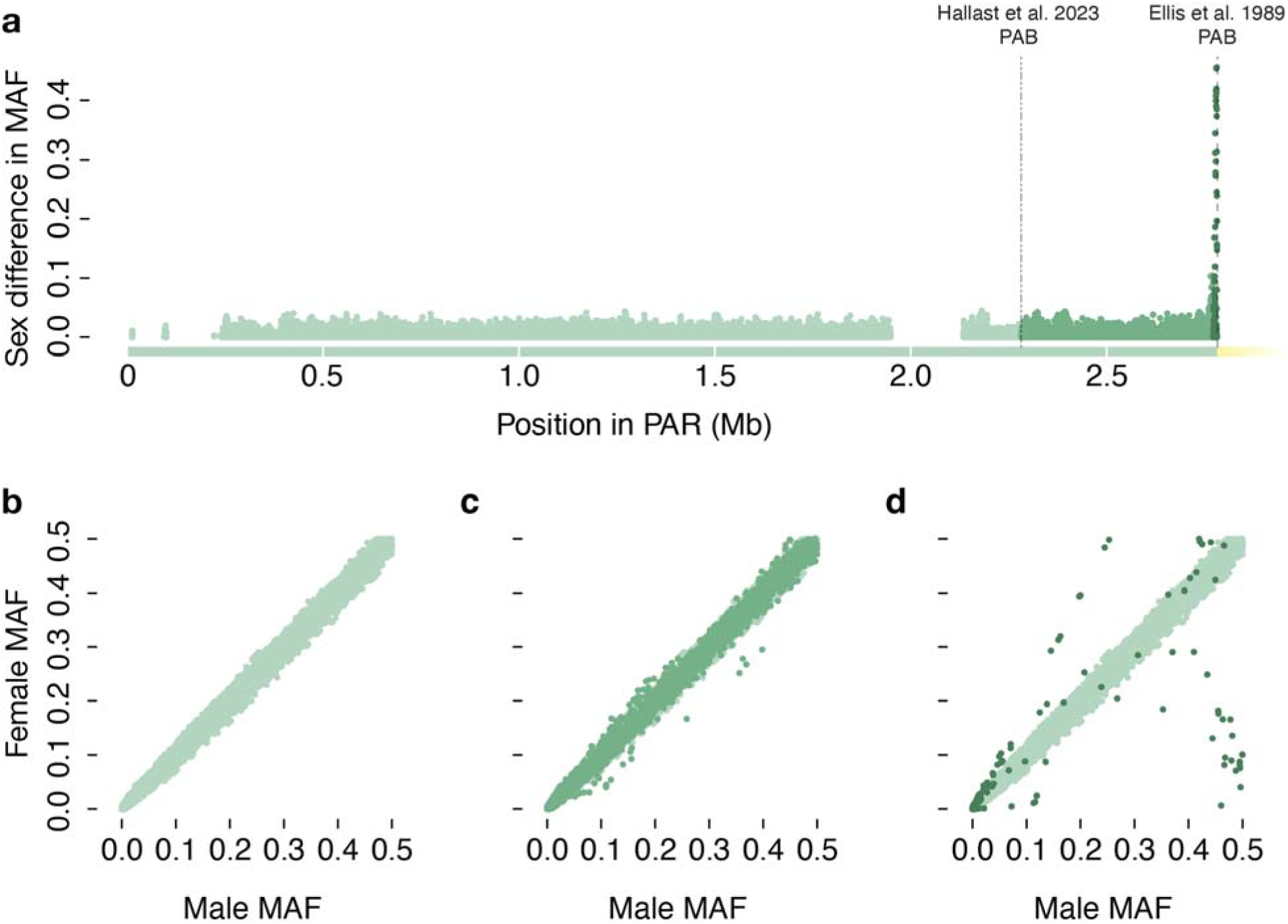
Sex differences in MAF only occur within 10 kb of the 1989 PAB. Minor allele frequency (MAF) calculated from 1000 genomes data for the human PAR. Dark green, sites <10 kb from the 1989 PAB; medium green, <500 kb from the 1989 PAB; light green, all other sites. (a) Absolute sex difference in MAF plotted against hg38 location. 2023 and 1989 PAB indicated by dashed lines. (b-d) Male versus female MAF; (b) tight correlation between male and female MAFs distal to the 2023 PAB; (c) similar correlation at most positions between the 2023 and 1989 boundaries; except (d) in the 10 kb immediately distal to the 1989 PAB.

### Single sperm sequencing

To capture ongoing crossover events in PAR1 from living individuals, we analyzed a massive single-sperm sequencing dataset (dbGaP project phs001887), containing Illumina reads from 31,228 sperm across 20 donors, which has not been previously analyzed for PAR1 crossovers^32^. We identified X– and Y-bearing sperm from the depth of reads aligning to the NPX and NPY and used this information to phase variant sites in PAR 1. Across all donors, we observed 795 sperm with unambiguous crossovers between the 1989 and 2023 PABs (Supplemental Table 3). We calculated the recombination rate between all genotyped variants in PAR1 (Figure 5a, Supplemental Figure 1), and estimate the total genetic distance between the 1989 PAB and 2023 PAB to be 4.78 cM (Figure 5b). We observe a local increase in the recombination rate close to the 1989 PAB (Figure 5a), parallel what Hinch and colleagues observed using a pedigree-based genetic map^20^. We note that this local pattern of increased recombination near the 1989 PAB is not present across all donors (Supplemental Figure 1), consistent with the observation that PAR1 hotspots are rapidly evolving and differ among human populations^20^.

**Figure 5:**
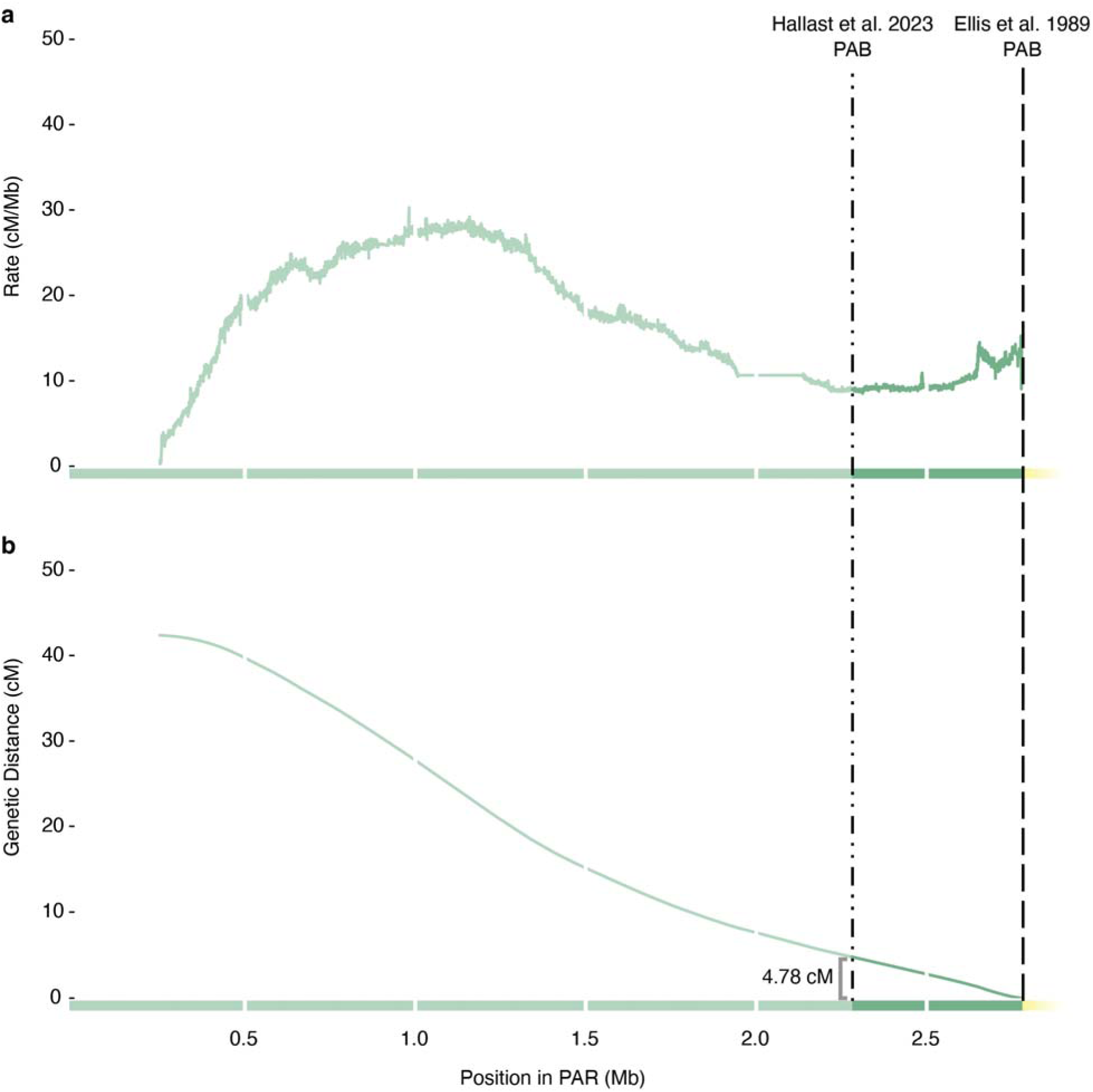
Single-sperm sequencing reveals crossovers between 2023 and 1989 PABs. Summary of Sperm-seq results across 20 donors showing (a) average local recombination rate between genotyped markers within PAR1, and (b) total genetic distance from the 1989 PAB. Medium green, region between 2023 PAB and 1989 PAB; light green, all other sites; white tickmarks every 500 kb.

### Reanalysis of Y chromosomes sequenced by Hallast and colleagues

All three of these fully independent analyses confirmed the location of the 1989 PAB, so we asked whether the sequences generated by Hallast and colleagues^1^ might also provide evidence for crossing-over in the 500 kb distal to the 1989 PAB. Of the 43 individual human male genomes assembled by Hallast and colleagues, ten have contiguous PAR1 assemblies, supported by high-quality differences between long reads originating from the X and Y chromosomes^1^. We restricted our analyses to these ten genomes where the provenance of the sequences within the PAR are unambiguous.

During human evolution, substitutions are predicted to accumulate on the NPY in a clock-like fashion, with no recombination. However, substitutions that arise in the PAR can be transferred from Y chromosomes to X chromosomes, or from one Y chromosome haplogroup to another through an X-chromosome intermediate, across generations. As a result, for sequences within the PAR, branch lengths calculated using the NPY haplogroup tree will show deviations from clock-like behavior. We constructed a multiple alignment of all ten contiguous Y-linked PARs (Supplemental Data 2). Using the phylogenetic relationships between these sequences based on their NPY haplogroup^1^, we conducted a likelihood ratio test (LRT) in 20-kb sliding windows of the alignment, comparing a single-clock model where all branches accumulate substitutions at the same rate, versus a local-clock alternative where rates are allowed to vary among branches (Figure 6, Supplemental Table 4). We observe a crisp change in the results of this test that coincides with the 1989 PAB (Figure 6). Windows proximal to the 1989 PAB are almost universally consistent with clock-like evolution along the haplogroup tree, as expected for NPY sequences. The rare exceptions may represent X-Y gene conversion events in homologous sequences. Distal to the 1989 PAB, the single-clock model can be rejected in the majority of windows. This can readily be explained by a history of crossing-over distal to the 1989 PAB, so that the Y-haplogroup tree does not accurately describe the relationship between these sequences.

**Figure 6:**
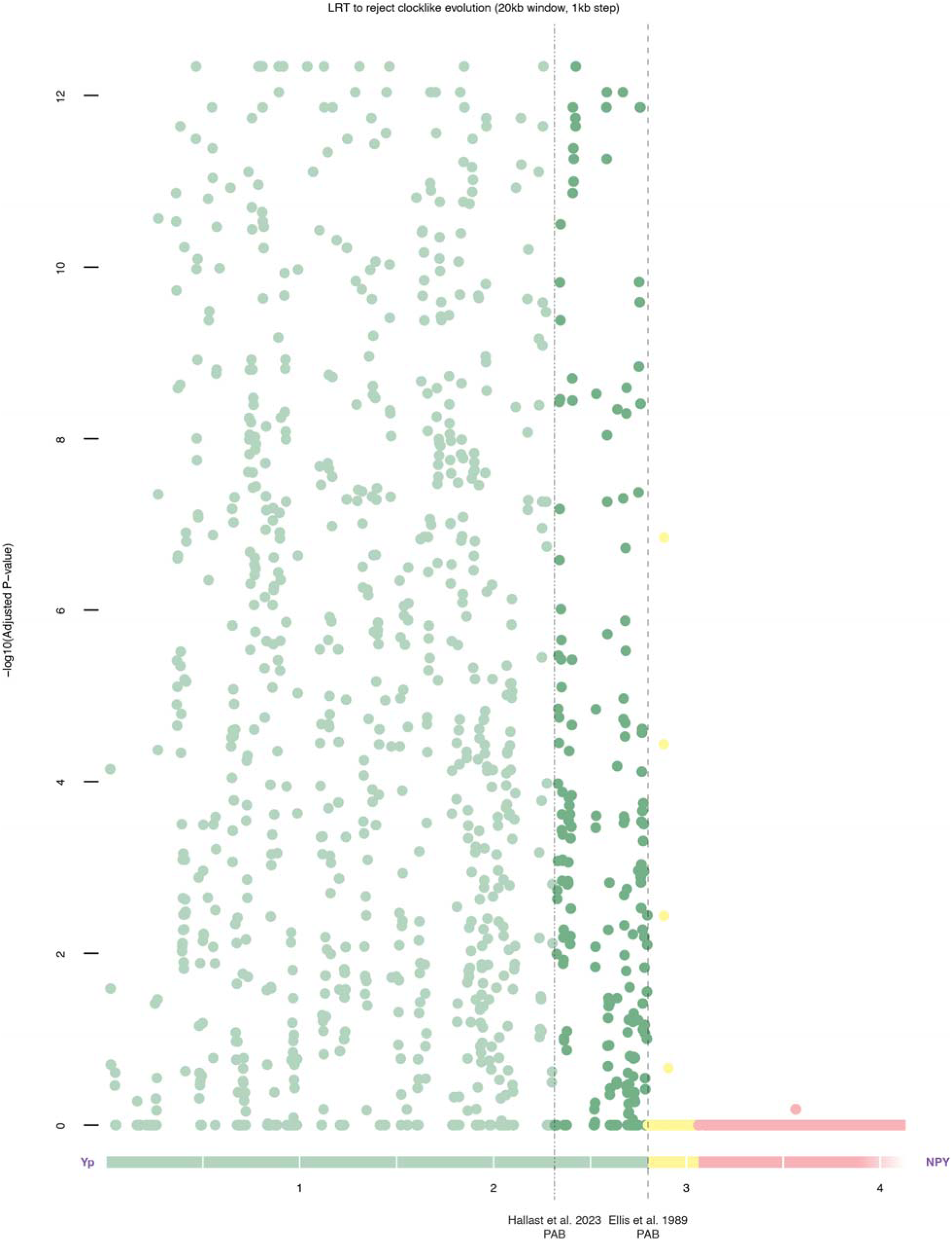
Y chromosome sequences distal to the 1989 PAB are inconsistent with phylogenetic tree of human NPY haplotypes. Results of LRT comparing single-clock versus local-clock models of evolution on the NPY haplogroup trees in 20-kb sliding windows across multiple alignment of ten contiguous Y chromosome sequences produced by Hallast and colleagues. Bonferroni corrected p-values for LRT plotted versus hg38 position. 2023 and 1989 PAB indicated by dashed lines. Windows distal to the 1989 PAB repeatedly reject the single-clock model.

Encouraged by these results, we further examined the homologous X and Y sequences from these ten genomes to ask how close to the 1989 PAB we could observe evidence of crossing-over. Following the procedure of Ellis and colleagues, we constructed a multiple alignment of these ten diverse X– and Y-linked sequences, along with sequence from a BAC clone from the Y chromosome spanning the 1989 PAB (Figure 7, Supplemental Data 3), and looked for polymorphisms shared by X and Y chromosomes. We observe the same fixed differences between human X and Y chromosomes at positions 41 and 45 bases distal to the Alu insertion that Ellis and colleagues described more than 30 years ago (Figure 7). Similarly, we see that all Y chromosomes share a haplotype commonly found on X chromosomes up to 245 bases distal to the Alu insertion (Figure 7). We observe five previously reported polymorphic sites (rs147962110, rs189310269, rs180883975, rs2534635, and rs5982851) where reference and alternate alleles are shared by X and Y chromosomes (Figure 7). The closest site, rs5982851, is only 246 bases distal to the Alu insertion that marks the PAB (Figure 7), while the next closest, rs2534635, was previously observed by Ellis and colleagues^16^. Thus, rather than overturning the 1989 PAB, the sequences produced by Hallast and colleagues reveal a historical recombination event 28 bases closer to the 1989 PAB than Ellis and colleagues initially observed, narrowing the possible location of the PAB to the 201-base interval between rs5982851 and the first fixed difference between the X and Y chromosomes.

**Figure 7:**
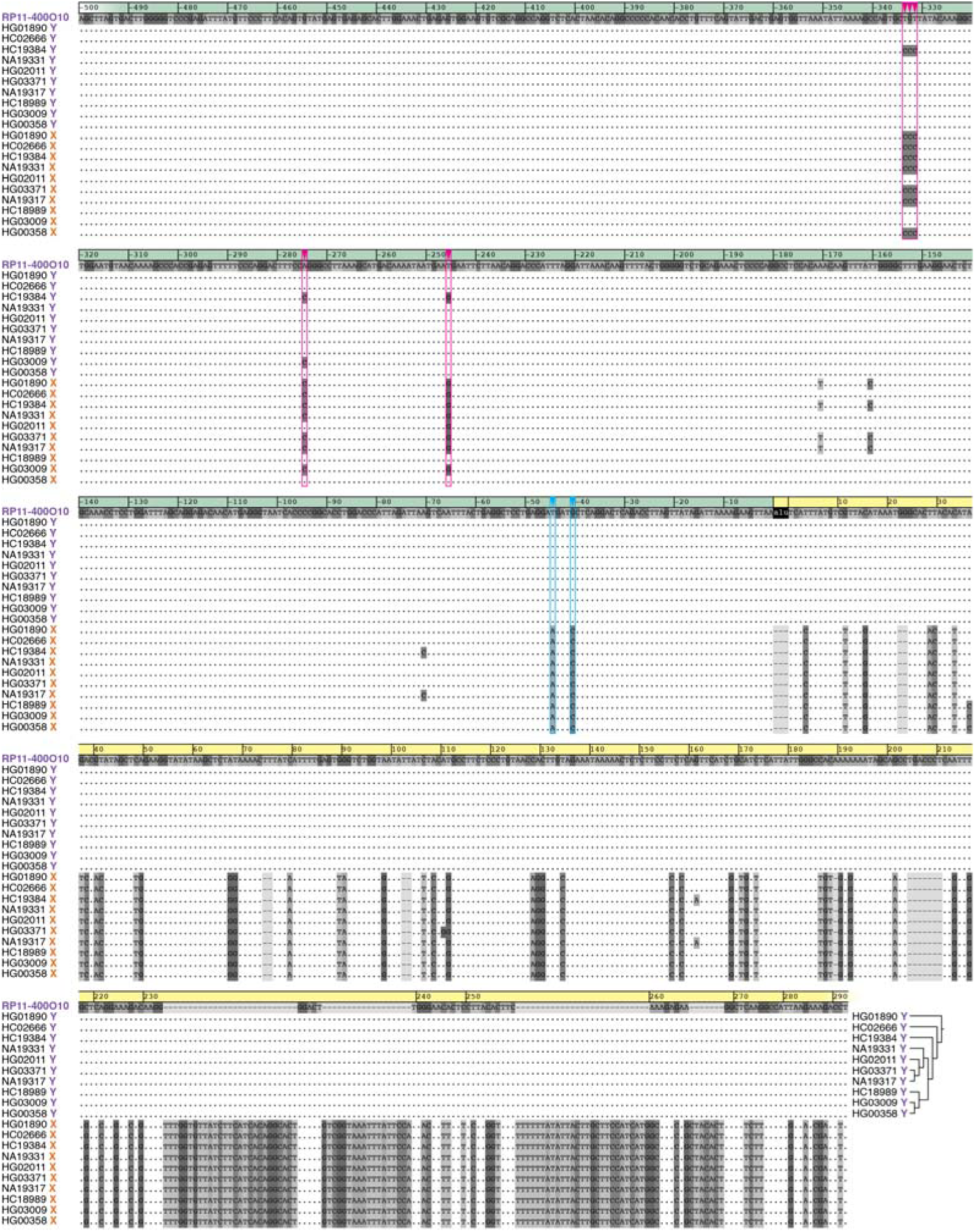
Polymorphisms shared by X and Y chromosomes within 500 bases of the 1989 PAB. Multiple alignment of human sex chromosome sequences showing the 1989 PAB at or near the Y-specific Alu insertion (black). Dots indicate identity with RP11-400O10 clone from human Y; coordinates relative to Alu insertion. Tree indicates NPY haplogroup tree for ten contiguous Y chromosome sequences produced by Hallast and colleagues. Magenta arrowheads and boxes mark polymorphisms shared by X and Y; cyan arrowheads and boxes mark fixed differences.

## Discussion

Our present understanding of the biology of the sex chromosomes has been informed by close study of crossing-over between the X and Y chromosomes. During meiosis, crossing-over generates a physical link between each pair of homologous chromosomes, providing the tension required to ensure that homologs can be segregated faithfully. The PARs are responsible for providing this obligate crossover between the X and Y chromosomes during male meiosis. Without this crossover, meiosis generates XY sperm that produce 47,XXY offspring that develop Klinefelter syndrome. In contrast, when there are crossovers involving the NPX or NPY, sperm will carry structurally anomalous X and Y chromosomes that produce 46,XX male (DSD) and 46,XY female (DSD) offspring. All three conditions result in infertility, imposing strong selection to maintain the PAR. The highly differentiated human X and Y chromosomes evolved from what were once an ordinary pair of autosomes that crossed over freely. This divergence was the result of a series of rearrangements over the last 200 million years that first suppressed crossing-over around *SRY*, the testis-determining gene, initially creating a PAB, and then repeatedly shifted the PAB to expand the NPY at the expense of the PAR^40^. The absence of crossing-over in males radically altered both the NPX and NPY; 97% of ancestral NPY genes were lost to genetic decay, while their counterparts on the NPX evolved new regulatory mechanisms to maintain the ancestral gene dose in both sexes^41^. As a result, the PAB forms the frontier between the sex-specific biology of the NPX and NPY, and the sex-shared biology of the PAR. As we continue to investigate the genetics of sex differences, the PAB remains a critical landmark on the human sex chromosomes.

For 40 years the PAB has been defined by the pattern of crossing-over. The available evidence overwhelmingly supports the 1989 PAB in PAR1, rather than a site 500 kb more distal. Crossing-over between the X and Y chromosomes has maintained high X-Y identity across the region distal to the 1989 PAB since the divergence of apes and Old World monkeys, around 30 million years ago^42^. Within the human population, crossovers occurred as close as 246 bases to the 1989 boundary, as inferred from alleles at polymorphic sites shared by both X and Y chromosomes. Human pedigrees and single-sperm typing data record ongoing crossovers throughout the PAR, including the 500 kb adjacent to the 1989 boundary. It is therefore clear that that the absence of structural variation cannot be used as a proxy for the absence of crossing-over. However, we note that our findings do not otherwise challenge the methods or findings of Hallast and colleagues, which in other respects agree closely with previous characterizations of the human Y chromosome.

## Acknowledgments

This work was supported by the Whitehead Institute, the Howard Hughes Medical Institute, the Simons Foundation Autism Research Initiative, and generous gifts from The Brit Jepson d’Arbeloff Center on Women’s Health, Arthur W. and Carol Tobin Brill, Matthew Brill, Charles Ellis, the Brett Barakett Foundation, the Howard P. Colhoun Family Foundation, the Seedlings Foundation, and the Knobloch Family Foundation.

## Author Contributions

Conceptualization: D.W.B., J.H., H.S., D.C.P.; Data curation, Formal Analysis, Methodology, Software, Validation: D.W.B., J.H., H.S., E.C.O.; Visualization: D.W.B., E.C.O.; Funding acquisition, Project administration, Supervision: D.C.P.; Writing – original draft: D.W.B.; Writing – review & editing: D.W.B., J.H., H.S., E.C.O., D.C.P.

## Data and Code Availability

This study did not generate datasets. Code for analysis of Sperm-seq data is available at: https://github.com/dwbellott/sperm_seq_scripts

## Figures and Legends

**Supplemental Figure 1:**
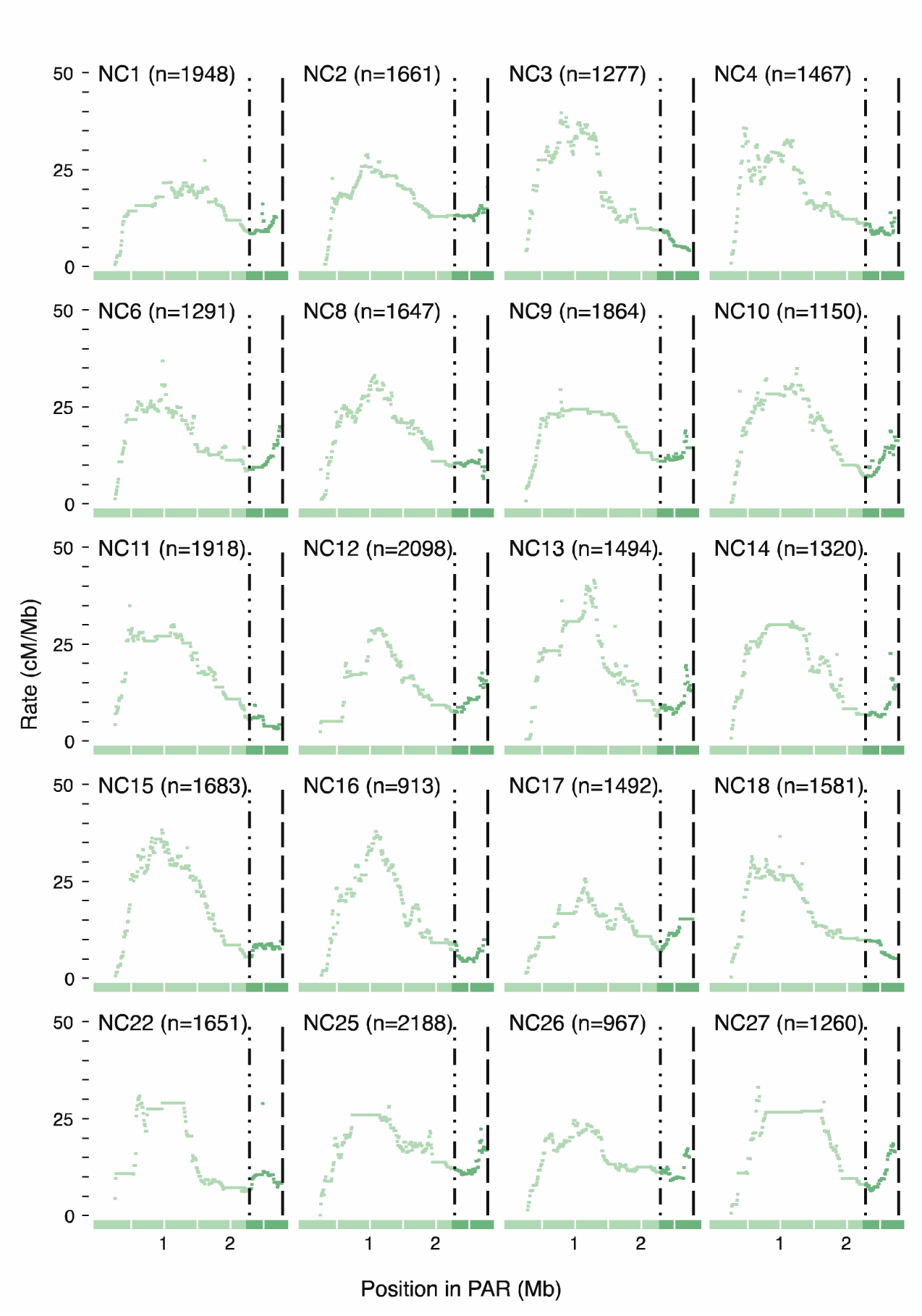
Single-sperm sequencing results. Summary of Sperm-seq results for 20 individual donors. Each plot shows local recombination rate between genotyped markers within PAR1; number of genotyped sperm indicated for each donor; medium green, region between the 2023 and 1989 PAB; light green, all other sites; white tickmarks every 500 kb.

