## Supplementary figures and images for "Where is the boundary of the human pseudoautosomal region?"

### Supplemental Figure 1

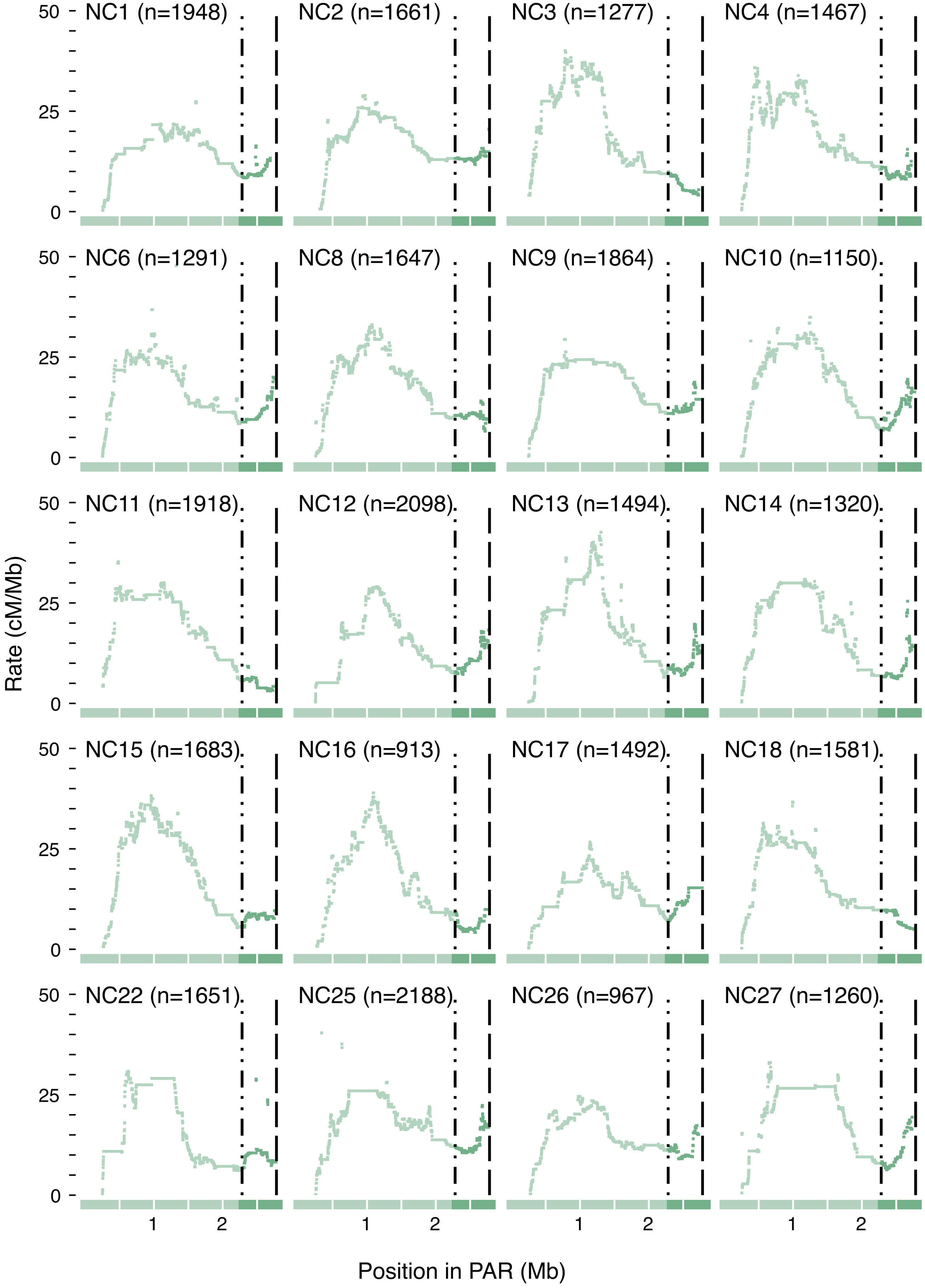
